# Transdermal patch containing physostigmine and procyclidine protects rhesus monkeys against VX intoxication

**DOI:** 10.1101/2020.10.16.343434

**Authors:** Young Jo Song

## Abstract

VX is an organophosphate cholinesterase inhibitor known as a chemical warfare agent. This study was designed 1) to determine the acute toxicity of VX in male rhesus monkeys by subcutaneous administration, 2) to evaluate the efficacy of a transdermal patch containing physostigmine and procyclidine. The median lethal dose (LD_50_) of the subcutaneous injection of VX was 15.409 ug/kg, which was calculated using the up-and-down dose selection procedure based on deaths occurring within 48 h. To test the efficacy of the transdermal patch, rhesus monkeys were treated with a patch (5×5 cm^2^) alone or in combination with post-exposure therapy comprising atropine plus 2-pralidoxime (2-PAM), and then administered subcutaneous injection of VX at various doses. The rhesus monkeys pretreated with the patch alone were 100% protected against 1.5×LD_50_ of VX, while the rhesus monkeys treated with the patch, atropine, and 2-PAM were 100% protected against 50×LD_50_ of VX. This study demonstrated that patch pretreatment in conjunction with atropine and 2-PAM treatment is an effective regimen against high doses of VX.

## Introduction

VX, an organophosphorus compound and nerve agent known for its use as a chemical warfare agent, is an extremely lethal chemical (Zurer, 1998). It is a potent and irreversible acetylcholinesterase (AchE) inhibitor. The high toxicity of VX contributes to its specific reaction with AchE. Unlike G agents, VX is more stable, more resistant to detoxification, less volatile, and penetrates skin more efficiently (U.S. Department of the Army, 1974). VX are attractive chemical weapons to terrorists because they are fast-acting even in minute quantities and can cause death by multiple routes, including inhalation (Munro, 1994). In the past, VX was used by the Aum Shinrikyo cult in Japan to kill people (Zurer, 1998). Recently, VX was used for an assassination in Malaysia in 2017 (Nakagawa and Tu, 2018).

Current antidotes for VX intoxication are composed of a combination of pretreatment with the reversible AChE inhibitor pyridostigmine (PYS) and post-exposure therapy with a three-drug regimen consisting of atropine sulfate, 2-pralidoxime (2-PAM), and diazepam (Dunn and Sidell, 1989; Shih et al., 2003; Vrdoljak et al., 2006). Although PYS effectively inhibits the enzyme AchE in a reversible manner, PYS can hardly penetrate the blood-brain barrier (BBB). Therefore, PYS only acts on the peripheral nervous system (Rickett et al., 1987). VX easily passes into the BBB because of its high lipophilicity, and both the central and peripheral nerves are severely damaged (U.S. Department of the Army, 1974). Recently, physostigmine (PHS) has been proposed as an alternative to PYS (Solana et al., 1990). Previous studies have reported that PHS is very effective against sarin or soman poisoning (Cho et al., 2012; Philippens et al., 2007). However, PHS has a short half-life in plasma and a narrow therapeutic efficacy, thus requiring administration as a sustained release formulation (Jenner et al., 1994). Nevertheless, the antidotal efficacies of PHS augmented synergistically when combined with procyclidine (PC) or scopolamine (Myhrer et al., 2010; Myhrer et al., 2004; Choi et al., 2004). Procyclidine is an antagonist of cholinergic and NMDA receptors (McDonough Jr. and Shih, 1995). It functions as an anti-convulsant and neuroprotective (Kim et al., 1997).

Our group has studied a transdermal patch system on a long-term basis to find better alternatives to nerve agents. In a previous study, we developed a transdermal patch containing PHS and PC and demonstrated that a prototype of this transdermal patch system fully protected rats, beagle dogs, and rhesus monkeys against soman poisoning (Cho et al., 2012; Choi et al., 2004; Kim et al., 2005). In addition, the patch system showed attenuation of clinical signs and seizures induced by soman toxicity. These effects led to the minimization of brain injury.

In this study, we first analyzed the drug concentration in the blood after applying patches on rhesus monkeys. Next, we evaluated the median lethal dose (LD_50_) of VX by subcutaneous injection in rhesus monkeys. To the best of our knowledge, this experiment has not been performed on rhesus monkeys before. Third, the effectiveness of the patch, alone or in combination with an antidote consisting of atropine plus 2-PAM were evaluated based on survival rate and clinical signs against VX intoxication in the rhesus monkeys.

## MATERIALS AND METHODS

### Chemicals

Atropine sulfate and 2-pralidoxime chloride were purchased from Sigma Aldrich (Sigma-Aldrich, St. Louis, MO). For animal injection, atropine (0.5mg/kg) and 2-pralidoxime (15mg/kg), based on the field formulas (Field Manual, FM 8-285) were dissolved in 0.9% saline and administered in a volume of 0.1ml/kg. VX of 98% purity were synthesized in single small scale facility of Agency for Defense Development, Republic of Korea. Stock solutions (2.0 mg/ml) of VX were prepared in iso-propanol and subsequent dilutions were performed in 0.9% saline for injection immediately before administration to animals.

### Transdermal patch

Matrix-type transdermal patch was made as previously described (10). In briefly, for the preparation of patch, desalted procyclidine (8.7mg) and physostigmine (34.8mg) in 7×7 cm^2^ were dissolved in ethyl acetate-ethanol and mixed with an acrylic adhesive DURO-TAK 387-2287/87-287 (National Starch and Chemical Co., USA).

### Animals

The fifty six male rhesus monkeys (*Macaca mulata*) were obtained from China by Woo Jeong Bio (Woo Jeong Bio, Suwon, South Korea). These monkeys were quarantined on arrival for one month and monitored for health condition. The animals were 3-4 years old and their body weights ranged between 3.0 and 4.5 kg. The animals were individually housed in stainless cages and maintained in a temperature (23+-2°C) and relative humidity (55+-3%) controlled vivarium under a 12h light-dark cycle. Water was available ad libitum and all monkeys are fed commercial primate pellets and fresh fruit. The Animal Care and Use Committee at the Agency for Defense Development approved the rhesus monkey experiments (ADD-IACUC-17, 18, 19).

### Drug concentration and Measurement of AchE activity in blood

To obtain profiles of blood concentration of PHS, PC and AchE inhibition rates, various size (3×4cm^2^, 5×5cm^2^, 7×7cm^2^) of patch were attached to the dorsal area of rhesus monkey after removal of the hair with clipper and a depilation cream (Table 2). Blood was collected with heparinzied tube at 0, 2, 4, 6, 8, 12, 24, 48, 72, 96 and 144 hours. AchE activity in blood was determined using modified Ellman method (21).

### LD_50_ of VX

Five monkeys were given doses of VX ranging from 20 to 12.6 μg/kg by subcutaneous (SC) challenge (Table 1). The experiments were performed using a sequential up-and-down method for small scales (19)Using this approach, a first animal was injected at specific dose and the outcome obtained before the next animal was challenged. The starting dose was 20 μg/kg, which is approximately halfway between the reported VX intravenous administration LD_50_ for rhesus monkeys (20). The dose of VX was usually increased if an animal lived and decreased if an animal died within 48hour. An estimate of the LD_50_ and confidence intervals were calculated by up-and-down method.

**Table 1.**
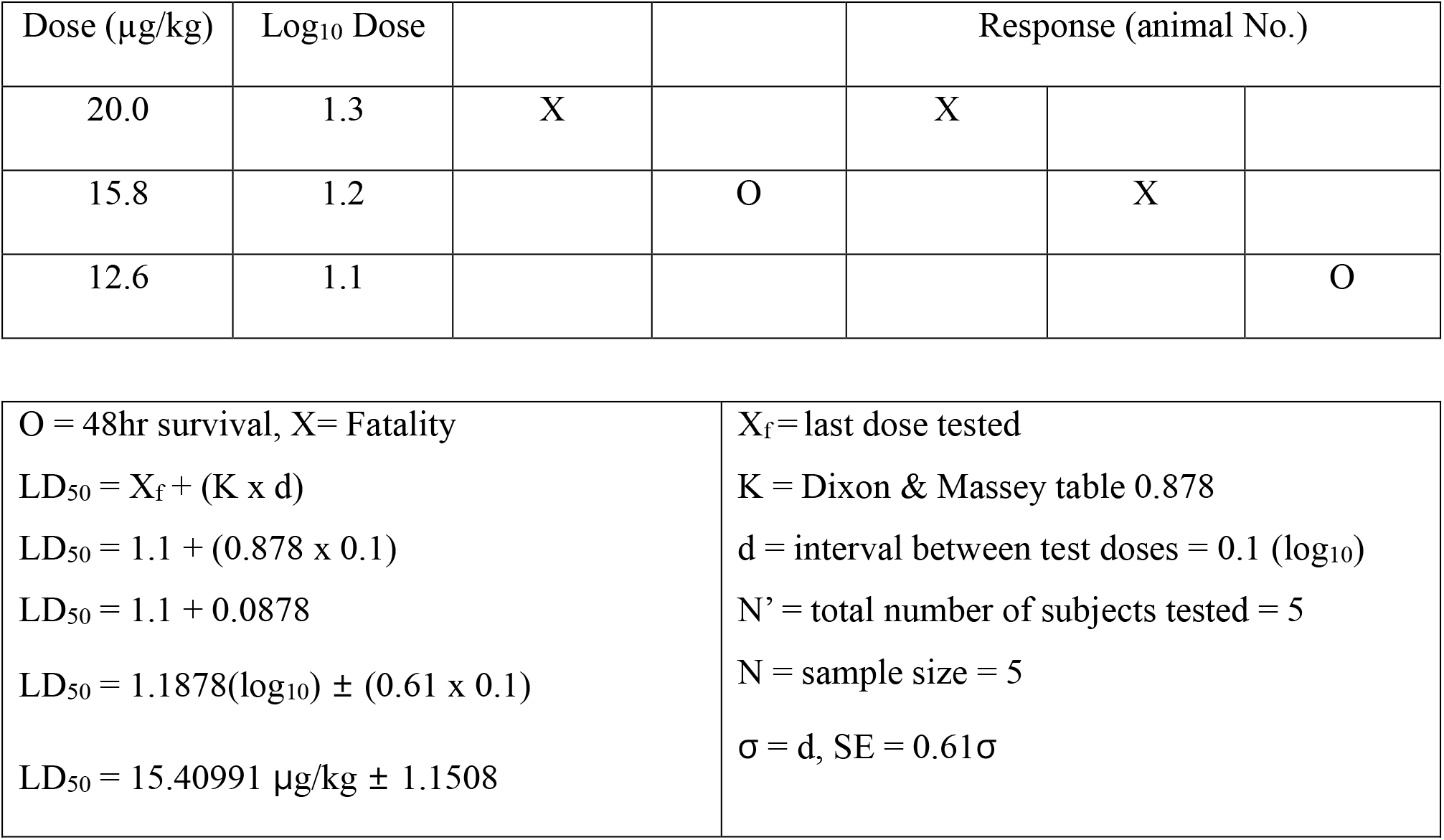
Summary of animal responses and LD_50_ calculation for the Up and Down method.

| Dose (µg/kg) | Log<sub>10</sub> Dose |  |  | Response (animal No.) |  |  |
| --- | --- | --- | --- | --- | --- | --- |
| 20.0 | 1.3 | X |  | X |  |  |
| 15.8 | 1.2 |  | O |  | X |  |
| 12.6 | 1.1 |  |  |  |  | O |

|  |  |
| --- | --- |
| O = 48hr survival, X= Fatality<br>$LD_{50} = X_f + (K \times d)$<br>$LD_{50} = 1.1 + (0.878 \times 0.1)$<br>$LD_{50} = 1.1 + 0.0878$<br>$LD_{50} = 1.1878(\log_{10}) \pm (0.61 \times 0.1)$<br>$LD_{50} = 15.40991 \mu\text{g/kg} \pm 1.1508$ | $X_f$ = last dose tested<br>K = Dixon & Massey table 0.878<br>d = interval between test doses = 0.1 (log<sub>10</sub>)<br>N' = total number of subjects tested = 5<br>N = sample size = 5<br>$\sigma = d$, SE = 0.61σ |

**Table 2.** Group of drug concentration and Ach E activity in the Rhesus monkey.

| Patch size | Animal No. | Blood collection |
| --- | --- | --- |
| 3×4 cm<sup>2</sup> patch | 5 | 0, 2, 4, 8, 12, 24, 48, 72, 96, 144 hr<br>(Total 10 times collection) |
| 5×5 cm<sup>2</sup> patch | 5 |  |
| 7×7 cm<sup>2</sup> patch | 5 |  |

### Efficacy of transdermal patch and combination therapy against VX toxicity

5×5 cm^2^ patch was attached on the dorsal area of the respective animal after removal of hair with a clipper and a depilation cream. The subjects were challenged with 2.0 ~ 1.5×LD_50_ VX by SC (Table 3). The challenge dose was calculated from the LD_50_ calculation experiment described above. For analyzing synergistic effect of combination therapy composed of transdermal patch and antidote, the monkeys, attached patch for 24 hours, were intramuscularly challenged with 10.0 ~ 100×LD_50_ of VX. In addition, the traditional antidote composed of atropine plus 2-PAM were treated intramuscularly within 1 minute after VX exposure (Table 4). The animals were monitored for cholinergic symptoms and survival.

**Table 3.** Effects of the patch against VX intoxication.

| Treatments | VX challenge Dose (LD<sub>50</sub>) | Animal No. | Survival No/Total No. (%) |
| --- | --- | --- | --- |
| None | 23.1 ug/kg (1.5LD<sub>50</sub>) | 1 | 0/1 (0) |
| 5×5 cm<sup>2</sup> patch | 23.1 ug/kg (1.5LD<sub>50</sub>) | 5 | 5/5 (100) |
| 5×5 cm<sup>2</sup> patch | 24.6 ug/kg (1.6 LD<sub>50</sub>) | 5 | 4/5 (80) |
| 5×5 cm<sup>2</sup> patch | 30.8 ug/kg (2.0 LD<sub>50</sub>) | 5 | 3/5 (60) |

**Table 4.** Effects of the combination therapy against VX intoxication.

| 5×5 cm<sup>2</sup> Patch | Atropine + 2PAM | VX challenge Dose (LD<sub>50</sub>) | Animal No. | Survival No /Total No |
| --- | --- | --- | --- | --- |
| X | O | 154.0 ug/kg (10.0LD<sub>50</sub>) | 1 | 1/1 |
| O | O |  | 1 | 1/1 |
| X | O | 462.0 ug/kg (30.0LD<sub>50</sub>) | 1 | 1/1 |
| O | O |  | 1 | 1/1 |
| X | O | 770.0 ug/kg (50.0LD<sub>50</sub>) | 6 | 3/6 |
| O | O |  | 6 | 6/6 |
| X | O | 1540.0 ug/kg (100.0LD<sub>50</sub>) | 2 | 0/2 |
| O | O |  | 2 | 1/2 |

## RESULTS

### Pharmacokinetic analysis of PHS and PC by application of transdermal patch

The drug pharmacokinetic parameters in the blood were summarized in the supplementary material. Pharmacokinetic parameters of PHS and PC in rhesus monkeys following application were evaluated at 0, 2, 4, 6, 8, 12, 24, 48, 72, 96 and 144 hours. The maximum concentration of PHS in the serum (C_max_) and the area under the concentration-time curve from zero time extrapolated to infinity (AUC_∞_) increased almost proportionally after patch application. The mean time to maximum concentration (T_max_) ranged from 5.2 to 12.0 h after patch application, then decreased gradually during the 3-day application. For the 3×4 and 5×5 patch, the drug levels showed similar kinetics in the blood, whereas the drug concentration of the 7×7 patch was higher than that of the patch during the 3-day application. Twenty-four hour application of the patch resulted in PHS and PC concentrations of 2.4–20.8 ng/ml and 45.3–115.0 ng/ml, respectively, in the blood. The PHS level returned to baseline three days after application, and the PC level returned to baseline three days after application.

### AchE inhibition in blood

Blood AchE inhibition reached its maximum level 6 h - 24h after patch application, which is consistent with the profile of PHS concentration. However, AchE inhibition rate was maintained at 26% for six days and slightly decreased in a time-dependent manner (Fig. 1).

**Fig. 1.**
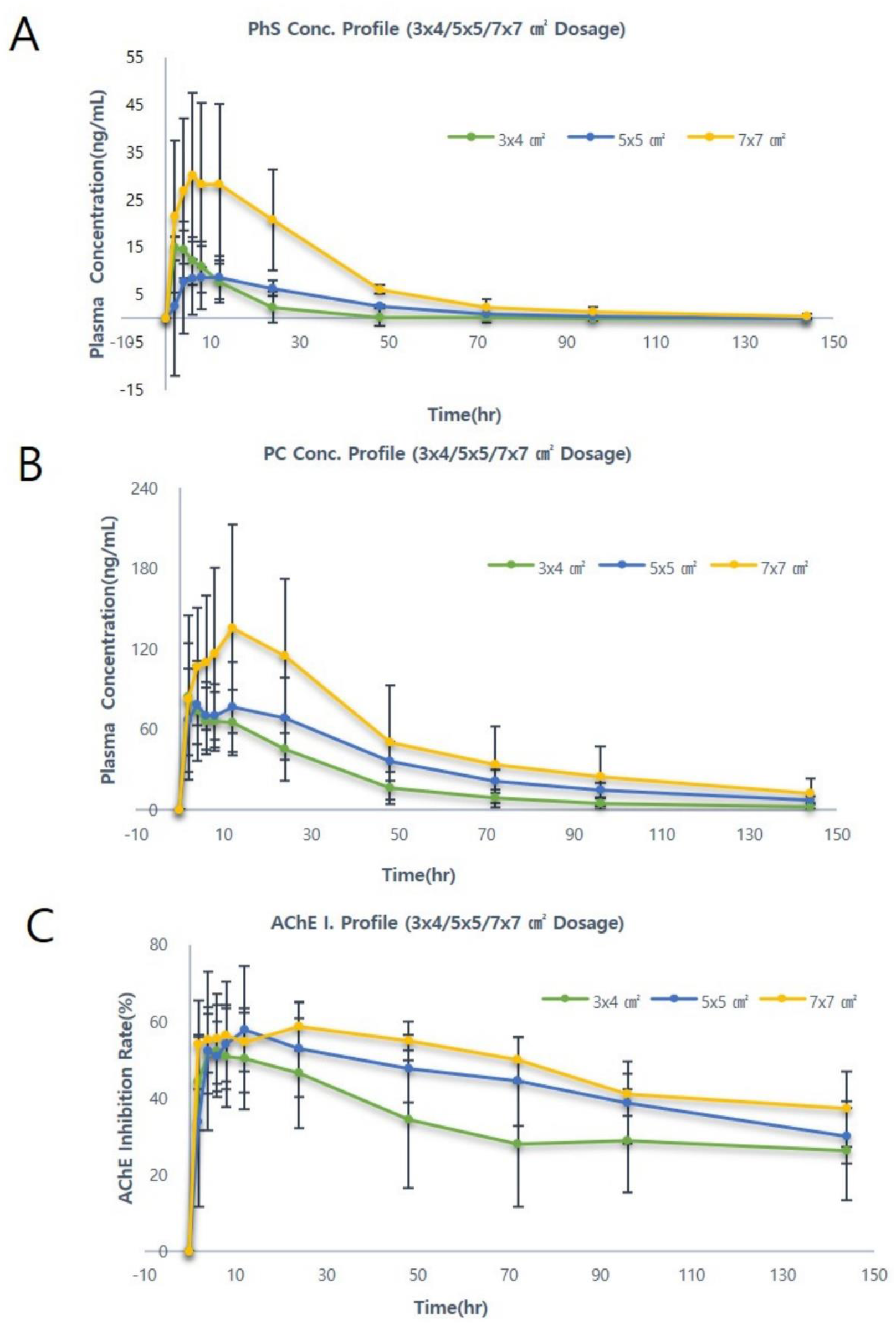
Analysis of blood concentration of physostigmine (A), procyclidine (B) and AchE inhibition rate (C) during attachment with patch.

### LD_50_ evaluation of VX

For LD_50_ evaluation of VX by subcutaneous administration in rhesus monkeys, a series of administrations was carried out according to the rule of increasing the dose following a survival and decreasing the dose following a death (Table 1). Within minutes of VX injection after a death, monkeys showed severe chewing and facial automatisms. This was followed by strong and continuous tremors throughout the body. This phase was followed by seizure, prostration, and unconsciousness. At the starting dose, death occurred within 24 h. At the second dose, the monkeys received 15.8 ug/kg of VX and exhibited delayed onset of seizures. After termination of the seizures, the animal slowly restored consciousness, although clinical signs persisted for 24 h. The surviving monkeys were able to sit in their cage 24 h after administration. The third dose had similar results as the first. At the fourth administration, animals dosed with 15.8 ug/kg of VX elicited severe signs of VX intoxication, and died within 24 h. At the fifth administration, animals survived at a dose of 12.6 ug/kg; however, they showed tremor, fasciculation, and salivation for 12 h after which normal behavior returned.

### Efficacy of transdermal patch

All animals exposed to VX were carefully monitored for clinical signs and mortality. The monkeys treated with patch for 24 h were exposed to various doses, and the results are summarized in Table 3. The patch group exposed to a 1.5×LD_50_ dose showed mild symptoms of intoxication, such as transient tremors and seizures on the day of VX challenge. Weak locomotor activity was observed after 48 h, and then returned to normal three days after VX challenge. Monkeys with patches, exposed to 2×LD_50_ dose (30.8 ug/kg) of VX exhibited salivation, tremors, and defecation followed by seizures, and three monkeys out of five died within 24 h. For further experiments on the combinational effects of the prophylactic patch and the therapeutic atropine and 2-PAM, patch-treated monkeys were additionally administered atropine and 2-PAM one minute after injecting various doses of VX. The results are summarized in Table 4. Despite the intoxicative symptoms induced by VX challenge, monkeys treated with patch plus antidote had 100% survival against 30×LD_50_ of VX. However, monkeys treated with antidote alone died within 24 h. Remarkably, monkeys treated with patch plus antidote had 50% survival against 100×LD_50_ of VX.

## DISCUSSION

Carbamates have demonstrated effective antidotal efficacy as anticholinergic drugs against nerve agents in guinea pigs and marmoset monkeys (Leadbeater et al., 1985; Philippens et al., 1998). Pretreatment with carbamates attenuates seizure and brain damage (Shin et al., 1992; McDonough Jr. and Shih, 1993). Therefore, our group has focused on developing a simple prophylactic regimen containing carbamates such as PHS. A previous study demonstrated that a combination of PHS and PC was effective in the prevention of lethality and seizure against organophosphorus intoxication (Kim et al., 2002). In addition, continuous infusion of PHS and PC by osmotic pump was found to be highly effective (Choi et al., 2004). More recently, a transdermal patch containing PHS and PC was developed, and its prophylactic efficacy against soman poisoning was demonstrated in rhesus monkeys and beagle dogs (Cho et al., 2012; Kim et al., 2005). In this study, we showed that a transdermal patch alone or in combination with traditional antidotes protects rhesus monkeys against VX. A transdermal patch alone containing PHS 34.8 mg and PC 8.7 mg protected rhesus monkey against 1.5×LD_50_ of VX. Furthermore, the treatment regimen composed of pretreatment with patch and post-exposure treatment with atropine plus 2-PAM exerted synergistic survival against 50×LD_50_ of VX. In contrast, monkeys that received only post-exposure therapy with atropine plus 2-PAM showed a 50% survival rate against 50×LD_50_ of VX. The surviving monkeys exhibited profound symptoms of intoxication, such as tremors and seizures. PHS plays a role in inhibiting AchE temporarily, thus shielding AchE from permanent inhibition by inflow of the nerve agent (Bartolini et al., 1973). In addition, PHS can protect the central nervous system because it is an unquaternized carbamate that penetrates the BBB (Solana et al., 1990). Therefore, PHS has been studied as a promising prophylactic agent against organophosphorus intoxication. In particular, when PHS was used with PC, synergistic effects were observed in marmoset monkeys and guinea pigs (Philippens et al., 2007).

PC is an antagonist of both nicotinic and muscarinic receptors. Therefore, it has antagonistic activity on NMDA receptors. Involuntary seizure after nerve agent intoxication stimulates NMDA signaling. Thus, treatment regimens including PHS (carbamate) combined with PC (anti-NMDA) would be synergistic against organophosphorus toxicity and brain damage after nerve agent intoxication. Our group performed brain histopathology to confirm whether the patch attenuated brain injury in a previous study. Severe neuropathological changes, such as eosinophilic neuronal injury, neuronal vacuolation, and shrunken neurons as a result of seizure were observed in the atropine plus 2-PAM-treated animal brains. In contrast, patch-treated animal brains showed normal features of histology (Cho et al., 2012). In this respect, the patch demonstrated brain protection against a nerve agent.

Rhesus monkeys have been used previously as a non-human primate research species of choice to evaluate nerve agent toxicity and efficacy of medical countermeasures. However, the supply of rhesus monkeys has been reduced recently, and the availability of animals has greatly decreased. The LD_50_ of VX administered through the subcutaneous, intramuscular, and intravenous routes in rhesus monkeys has not been published. Previous studies performed small-scale experiments using 2 or 3 monkeys and showed that rhesus monkeys treated with VX developed tremors, severe seizures, and death (Raveh et al., 1997). Therefore, researchers have predicted the LD_50_ dose of VX on the basis of the results of the following animals. The LD_50_ of VX in mice, rats, rabbits, and guinea pigs was valued at 9 to 16 ug/kg (Wolfe et al., 1987). The LD_50_ dose of VX in cats was reported at 5 ug/kg by intravenous route (Rickett et al., 1986). These differences were thought to be related to endogenous carboxylesterase in the body of animals (Maxwell et al., 1992). As a result, the LD_50_ of VX administered intravenously was estimated to be < 7.5 ug/kg. The present study demonstrates the LD_50_ of VX (15.4 ug/kg, sc) in rhesus monkeys for the first time. The subcutaneous LD_50_ dose was two times higher than that of the predicted intravenous LD_50_. In the soman agent, the intravenous LD_50_ dose was 5.5 ug/kg and the subcutaneous LD_50_ dose was 12.3–13.0 g/kg in rhesus monkeys. Therefore, the VX LD_50_ dose is considered to be similar to that of soman LD_50_ in rhesus monkeys. It should be noted that LD_50_ evaluation in this study using the up-down method was performed with fewer number of animals than with other methods such as probit analysis in previous studies. Notably, this result is considered an important administration route in the efficacy test of medical countermeasures for nerve agents. The present study also showed that the signs of VX intoxication in rhesus monkeys were consistent with what has been described in previous experiments. Within 20 min of VX injection, chewing and facial spasm were observed, followed by continuous tremor and convulsion within 30 min.

This study included a large-scale experiment using 60 monkeys and demonstrated that applying patches alone protected monkeys against 1.5×LD_50_ of VX, while combined administration with antidotes protected monkeys against 50×LD_50_ of VX. As a result, the preventive application of patches and the therapeutics of antidotes such as atropine and 2-PAM are very effective against VX. Recently, this transdermal patch cleared a nonclinical study and is currently under clinical trial.

## Acknowledgements

This work was supported by grant from Agency for Defense Development, Republic of Korea (571665-271987001, 671665-271987001, 771665-271987001).

## Abbreviations

LD_50_: median lethal dose
2-PAM: 2-pralidoxime
AchE: acetylcholinesterase
PYS: pyridostigmine
PHS: physostigmine
PC: procyclidine
BBB: blood-brain barrier
C_max_: maximum concentration of drug in the serum
AUC_∞_: area under the concentration-time curve from zero time extrapolated to infinity
T_max_: mean time to maximum concentration

---

**Supplementary Table 1.** Physostigmine pharmacokinetic mean data.

| Patch size | AUC<br>(last)ng·h/ml | AUC<br>(last)ng·h/ml | Cmax<br>(ng/ml) | Tmax (hr) | T1/2 (hr) |
| --- | --- | --- | --- | --- | --- |
| 3×4 cm<sup>2</sup> Patch | 218.5 | 231.8 | 16.9 | 5.2 | 11.30 |
| 5×5 cm<sup>2</sup> Patch | 348.6 | 355.5 | 9.8 | 12.0 | 20.6 |
| 7×7 cm<sup>2</sup> Patch | 994.4 | 1023.0 | 27.1 | 7.0 | 23.9 |

**Supplementary Table 2.** Procyclidine pharmacokinetic mean data.

| Patch size | AUC<br>(last)ng·h/ml | AUC<br>(last)ng·h/ml | Cmax<br>(ng/ml) | Tmax (hr) | T1/2 (hr) |
| --- | --- | --- | --- | --- | --- |
| 3×4 cm<sup>2</sup> Patch | 2833.5 | 3017.8 | 91.3 | 4.4 | 35.0 |
| 5×5 cm<sup>2</sup> Patch | 4839.3 | 5461.7 | 85.2 | 12.4 | 49.3 |
| 7×7 cm<sup>2</sup> Patch | 11081.7 | 12211.2 | 195.6 | 10.5 | 45.2 |

